# Visual Response Properties in the Three Layer Turtle Visual Cortex

**DOI:** 10.1101/131805

**Authors:** Mahmood S. Hoseini, Jeff Pobst, Nathaniel C. Wright, Wesley Clawson, Woodrow Shew, Ralf Wessel

## Abstract

Turtle dorsal cortex provides us with unique insights into cortical processing. It is known to share many features with the mammalian hippocampus and olfactory cortex as well as geniculo-cortical areas in stem amniotes from which mammals evolved. To this end, we have used data from extracellular recordings from microelectrode arrays to study spatial and temporal patterns of responses to visual stimuli as seen in both local field potential and action potentials. We discovered surprisingly large receptive fields, responsiveness to a broad range of stimuli, and high correlation between distant neural ensembles across recording array. Moreover, we found significant response variability regarding latency and strength in the presence of adaptation to both ongoing and visually evoked activity.

## Introduction

Understanding evolutionary origins of the mammalian cortico-thalamic system will infiltrate our knowledge. The study was launched through reptilian brains since they are simpler that their mammalian counterparts (Butler and Hodos, 2005; Naumann et al., 2015). Comparing cortical circuits between mammals and reptiles opens up a window into the role of different brain areas in cognitive processing (Naumann et al., 2015). Turtles are of particular interest for comparative studies because they probably bear the strongest resemblance to the Triassic cotylosaurs (stem amniotes) from which they and all modern mammals evolved (Romer and Parsons, 1977). Concomitant with comparative studies characterizing spatiotemporal features of visual responses helps us to uncover the underlying circuitry in mammalian cortex and, ultimately, understanding how the cortex processes sensory information.

The turtle dorsal cortex is a convergent zone for the visual, auditory, somatic, and other sensory systems (Gusel’nikov et al., 1972). Nonetheless, it has mostly been studied with respect to visual processing. There are several studies on the spiking response to diffuse flashes (Gusel’nikov et al., 1972; Kriegstein, 1987; Mancilla et al., 1998) and moving black dots (Gusel’nikov and Pivovarov, 1977) as well as the size and organization of the receptive field (RF) of spiking cells (Mazurskaya, 1973a). However, there have only been very few studies focusing on voltage sensitive dyes (Prechtl et al., 1997) or local field potential (LFP; Bass et al., 1983; Prechtl, 1994; Prechtl and Bullock, 1994; Prechtl et al., 2000; Luo et al., 2010). Nevertheless, there exist very little agreement about different features of response such as RF size, adaptation, and direction tuning (Gusel’nikov et al., 1972; Mazurskaya, 1973a; Boiko, 1980; Mulligan and Ulinski, 1990; Luo et al., 2010).

Here, we have revisited some of these questions. The clues at our disposal to explore response spatiotemporal properties are extracellular recordings from microelectrode arrays (MEA) that capture key integrative synaptic processes in neural networks. We have found large RFs that typically cover over half of the visual field with no indication of direction selectivity in line with previous study (Boiko, 1980). Investigating spatial structure of the receptive fields, we have discovered no clear or well-defined retinotopic map from the visual field (VF) to the cortex. Furthermore, our results demonstrate a broad range of response latencies and a strong adaptation to both ongoing and visually evoked activity.

## Materials and Methods

### Ex Vivo Eye-Attached Whole Brain Preparation

All procedures were approved by Washington University’s Animal Care and Use Committees and conform to the guidelines of the National Institutes of Health on the Care and Use of Laboratory Animals. Adult red-eared turtles (Trachemys scripta elegans, 150 – 200 *g* weight, 12 - 15 *cm* carapace length) were studied. Rapid decapitation was performed following anesthetization with Propofol (10 *mg*/*kg*) (Ziolo and Bertelsen, 2009). We then removed the brain with the right eye attached and proceeded to hemisect the eye.

To access the ventricular surface of the left visual cortex, we cut off ∼1 *mm* of the left olfactory bulb, which provided a hole to start a rostral-caudal cut through the medial cortex. This cut continued from the olfactory bulb into the natural separation of the medial cortex from the septum (about 1/3 of the cortex) and continued further along the same line for ∼1 - 2 *mm* into the caudal cortex. Finally, two cuts were made from the medial cortex edge towards the dorsal cortex. These two cuts were started at roughly 1/3 and 2/3 the rostral-caudal length of the cortex and were made as short as possible while still allowing the cortex to be unfolded and pinned flat. This length was usually ∼2 *mm*.

After making the cuts in the cortex, it was placed in the recording chamber on either a Sylgard or agar surface, and insect pins were used to pin the cortex flat. Our electrodes were then placed in the flattened cortex. The eye and brain were continuously perfused with artificial cerebrospinal fluid (in mM; 85 NaCl, 2 KCl, 2 MgCl_2_, 45 NaHCO_3_, 20 D glucose, and 3 CaCl_2_ bubbled with 95% O_2_ and 5% CO_2_), adjusted to pH 7.4 at room temperature. To perfuse the eye without obstructing the image we project onto the retina, a small wick was made from a Kimwipe. The wick connected an artificial cerebrospinal fluid (ACSF) feed located ∼1 *cm* above and to the side of the eye to the inside edge of the hemisected eye. If any brain tissue were large enough to extend above the surface of the ACSF (e.g., the right cortex or the optic tecta), it would be cover with a small piece of Kimwipe so that it would also stay in contact with ACSF. Recordings began 2-3 hours after anesthetization.

### Visual Stimulation

For the included studies, three methods of visual stimulation were used.

#### LED Stimulation

For LED stimulation, a red LED was connected to the output of a National Instruments BNC-2090 terminal block connected to a National Instruments PCI-6024E DAQ board. This output was controlled with a custom LabView program on a computer running Windows 7. The mean light intensity at the retina was 60 *mW*/*m*^2^.

#### Monitor/Mirror Stimulation

For monitor/mirror stimulation, a 19” LCD monitor (Samsung model Syncmaster T190, 1440x900 pixels, contrast ratio=20000:1, response time = 2 ms) displayed the stimuli. This image was reflected off a mirror located across room above the tissue, and focused on the retina with a lens placed above the tissue (Fig. 1a). The mean light intensity at the retina from the monitor was 20 *mW*/*m*^2^.

**Fig. 1.**
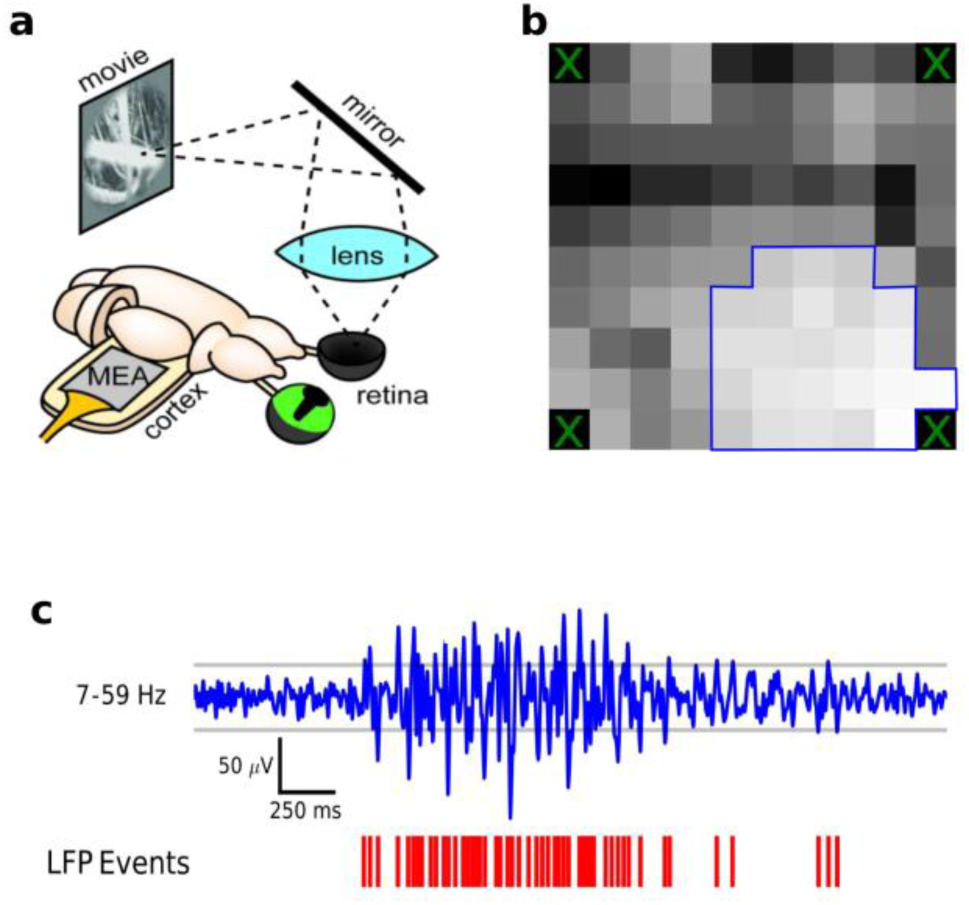
Experimental setup and quantifying LFP events. **a** Our experimental set up for experiments done with a monitor and mirror. The visual stimuli are presented on a monitor. The image reflects off a mirror and through a lens to form a picture on the retina of the turtle’s hemisected eye. The multielectrode array (MEA) is placed in the unfolded cortex. **b** Visual responsiveness across the MEA. Each square represents an electrode. The background color for each square indicates strength of their visual responses with black being the weakest and white being the strongest. Enclosed blue region indicates visual responses stronger than the threshold and being included in the analyses. **c** LFP events are referred to threshold crossings. A filtered LFP response is shown in blue with gray lines as ±3 standard deviation, and threshold crossings are marked in red as LFP events.

#### Projector Stimulation

For projector stimulation, a small projector was placed above the retina and focused with a system of lenses (Aaxa Technologies, P4X Pico Projector, 1440x900 pixels). The mean light intensity at the retina from the projector was 1 *mW*/*m*^2^. Both monitor/mirror and projector stimulation was provided using software written in python on a computer running Ubuntu 10.4. Visual stimuli included black dots moving on a white screen, naturalistic video, and red LED flashes.

### Data Acquisition

#### Microelectrode Array Recordings

Data were collected at a 30 *kHz* sampling rate using the Cerebus data acquisition system by Blackrock Microsystems. Two different styles of microelectrode arrays were used for our recordings. For some recordings, we used a 96-channel Utah array (Fig. 1b; 10x10 square grid, 400 *μm* inter-electrode spacing, 500 *μm* electrode length, no corner electrodes, Blackrock Microsystems). For others, we used an array of shank electrodes (4x4 array of shank electrodes with 8 recording sites on each electrode, 300 *μm* and 400 *μm* x and y distance between shanks and 100 *μm* between recording sites along a shank, Neuronexus). We attached either array to a post fastened to a micromanipulator (Sutter, MP-285). The Utah array was inserted to a depth of 250 - 500 *μm* starting from the ventricular side of the unfolded cortex such that the plane of electrodes was parallel to the dorsal surface of the cortex. The array of shank electrodes was inserted deep enough to span the entire depth of the cortex.

We recorded wideband (0.7 *Hz* – 15 *kHz*) extracellular voltages relative to a silver chloride pellet electrode in the tissue bath.

#### Single Electrode and Tetrode Recordings

For our experiments using single electrodes, we used tungsten electrodes (500 *k*Ω part #WE30030.5H5 from MicroProbes and 1000 *k*Ω catalog #573520 from AM Systems). For some experiments, we also used homemade tetrodes with resistances between 250 *k*Ω and 350 *k*Ω. These were made by twisting four 12.7 *μm* nickel chromium wires together, applying heat with a heat gun (Weller 6966C) and cutting the twisted wires at an angle to expose the ends for recording (Saha et al., 2013). Recordings were taken in reference to a silver chloride ground wire sitting in the bath.

The signals from these electrodes were recorded with an AM Systems Model 1800 amplifier connected to a National Instruments PCI-6024E 12-bit DAQ board through a National Instruments BNC-2090. The data were collected at 20 *kHz* using Labview.

### Filtering for Local Field Potential

To study LFP, it is useful to filter out other frequencies. For our LFP analysis, we used the PyWavelets package to perform wavelet filtering (Wiltschko et al., 2008). We used Daubechies wavelets with a minimum level of 9 and a maximum level of 11. For our 30 *kHz* data, this corresponds to a pass-band of ∼7 - 59 *Hz*.

*Defining an LFP Event.* There are many ways one can quantify the size of an LFP response. One method we use throughout this paper is to look for threshold crossings of the extracellular signal after filtering it to the frequencies we are interested in. As a threshold for LFP events we used 3 times standard deviation of the filtered signal (Fig. 1c). Therefore, when we refer to LFP event count, we are simply referring to a number of threshold crossings.

*Determining Visually Responsive Electrodes*. When recording from the 96 electrodes of the MEA, only a subset of electrodes would actually have a strong visual response. In order to get clean results, it is necessary to do our analysis on only that subset of electrodes.

To consistently and systematically determine which electrodes to include, we created an algorithm to test for visual responsiveness. Roughly speaking, an electrode was considered visually responsive if the typical level of activity following visual stimulation was sufficiently greater that spontaneous ongoing level activity. Specifically, we established the spontaneous ongoing level of activity, *a*_*ong*_, by taking the average activity from 4 *s* windows immediately preceding the presentation of the stimulus. Similarly, the amount of visually evoked activity, *a*_*evoked*_, was the average of the 4 *s* windows immediately following the onset of the visual stimulus. We then calculate the decrease in activity (how much lower is the ongoing activity level than the evoked activity level) as 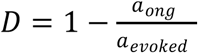. Finally, we classify an electrode as visually responsive if the decrease in activity is greater than 0.75 (Fig. 1b).

### Defining Receptive Field Similarity

To quantify RF similarity between nearby electrodes, we calculated the amount of overlap between the RFs from pairs of electrodes. Specifically, we binned the dots paths in visual field (bin size = 8 visual degrees), calculated the normalized average LFP response to stimulation in each of those bins, and for each electrode pair, we calculated the RF overlap (also called similarity) by summing the smaller of the two normalized response values (one for each electrode) over all bins.

The normalization of the average LFP response in each bin was done by dividing the average response by the sum of the average responses over all bins (or, in the case of direction specific RF similarity, by dividing by the sum of the average responses over only the bins for the angle of interest). Consequently, the sum of the normalized responses over all bins was always one, the minimum similarity between two electrodes was zero, and the maximum was one.

## RESULTS

### Overview

To quantify the spatiotemporal structure of visual responses we recorded extracellular neural activity using MEAs inserted into the geniculo-recipient dorsal cortex of the turtle eye-attached whole-brain *ex vivo* preparation during visual stimulation of the retina (Fig. 1a). First, we explore spatial properties in terms of RF size, RF similarity, and direction sensitivity; and then temporal features are quantified in terms of response duration, latency, and adaptation.

### Spatial Properties

LFP traces were recorded in response to black dots moving across a white background (8 degrees diameter dot moving at 40 *deg*/*s*). Dots move in several different directions (either 4 or 8 different angles), and for each angle, dots move along 8 different straight paths spanning the visual field. Our probe of the spatial structure includes RF size, RF similarity, and direction selectivity and is as follows.

### 1. RF size

LFP in response to moving dots on the screen is used to determine the size of RF. For each moving dot along a path, we mark LFP events of a given electrode (Fig. 1c; see Materials and Methods). In order to aid visualization, LFP events are plotted somewhat offset from the actual path of the presented dots. Events are color coded according to the direction of the moving dots. Responses in each direction over different trials and paths are summarized in the distribution of the average LFP event count with the trial-to-trial variability shown as the filled region around the average line (Fig. 2).

**Fig. 2.**
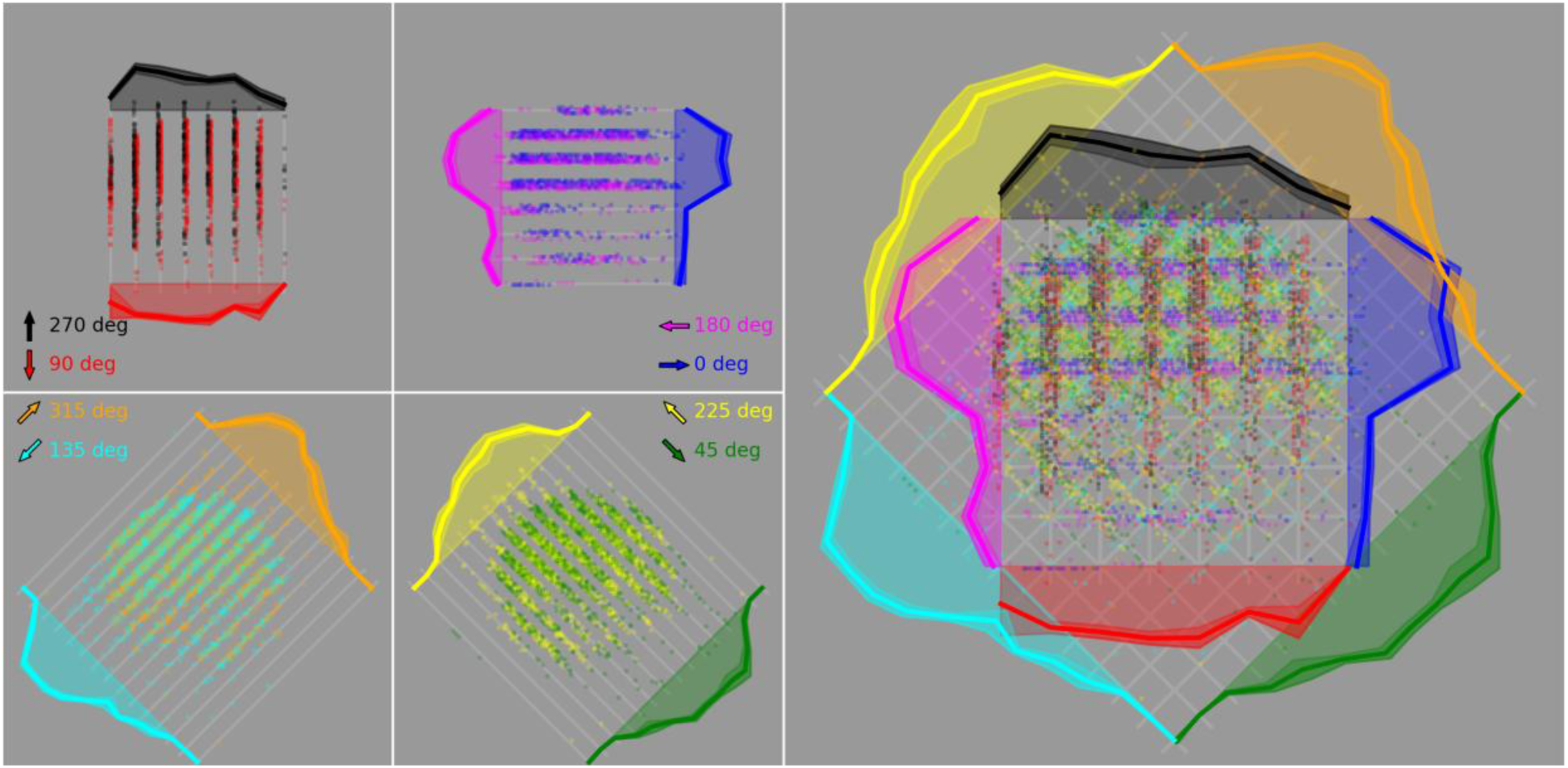
RF of the LFP signals covers over half of the visual field. Each grey square represents the visual field. Dots move along 8 different paths represented by light grey lines in 8 different directions. LFP events are color coded according to the direction of moving dot. In order to aid with visualization, the LFP events are plotted somewhat offset across each path. The solid lines show the across-trial average of LFP event count with the standard deviation shown as the filled region around the average solid line. On the left we presented the data from one electrode only for dots moving in opposite directions with respect to turtle’s visual streak (4 subplots for total of 8 directions). The figure on the right shows all 8 directions together. This shows that this electrode is responding to stimuli presented across a large region of the visual field. Retina is briefly exposed to the dots moving along the first and last path and, therefore, LFP responses are diminished. On the other hand dots presented in the middle paths usually evoke large responses.

Typically, near the edges (for the first and last paths) LFP response diminishes. This is due to the fact that for those paths dots spend less time moving in the visual field (Fig. 2). More importantly, the RF of the LFP seems to span large areas of the visual field in different preparations (Fig. 3) as well as across different electrodes of MEA (Fig. 4). We see that RFs commonly cover over half of the visual field.

**Fig. 3.**
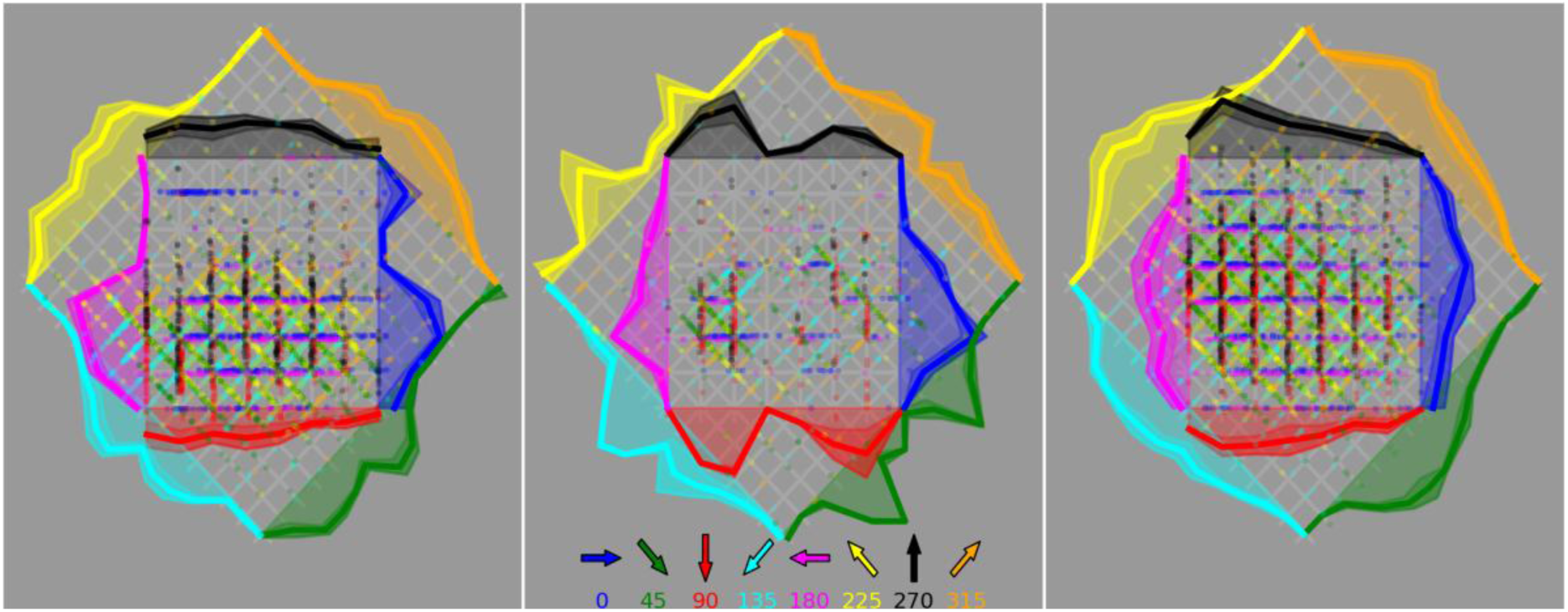
RFs are considerably large for in three different preparations. Responses to black dots moving in different directions on a white screen from the LFP of a single electrode in three different preparations exhibit RFs that cover a considerably large region of the visual field. The middle figure shows that binomial distribution is possible in which middle paths do not evoke strong responses.

**Fig. 4.**
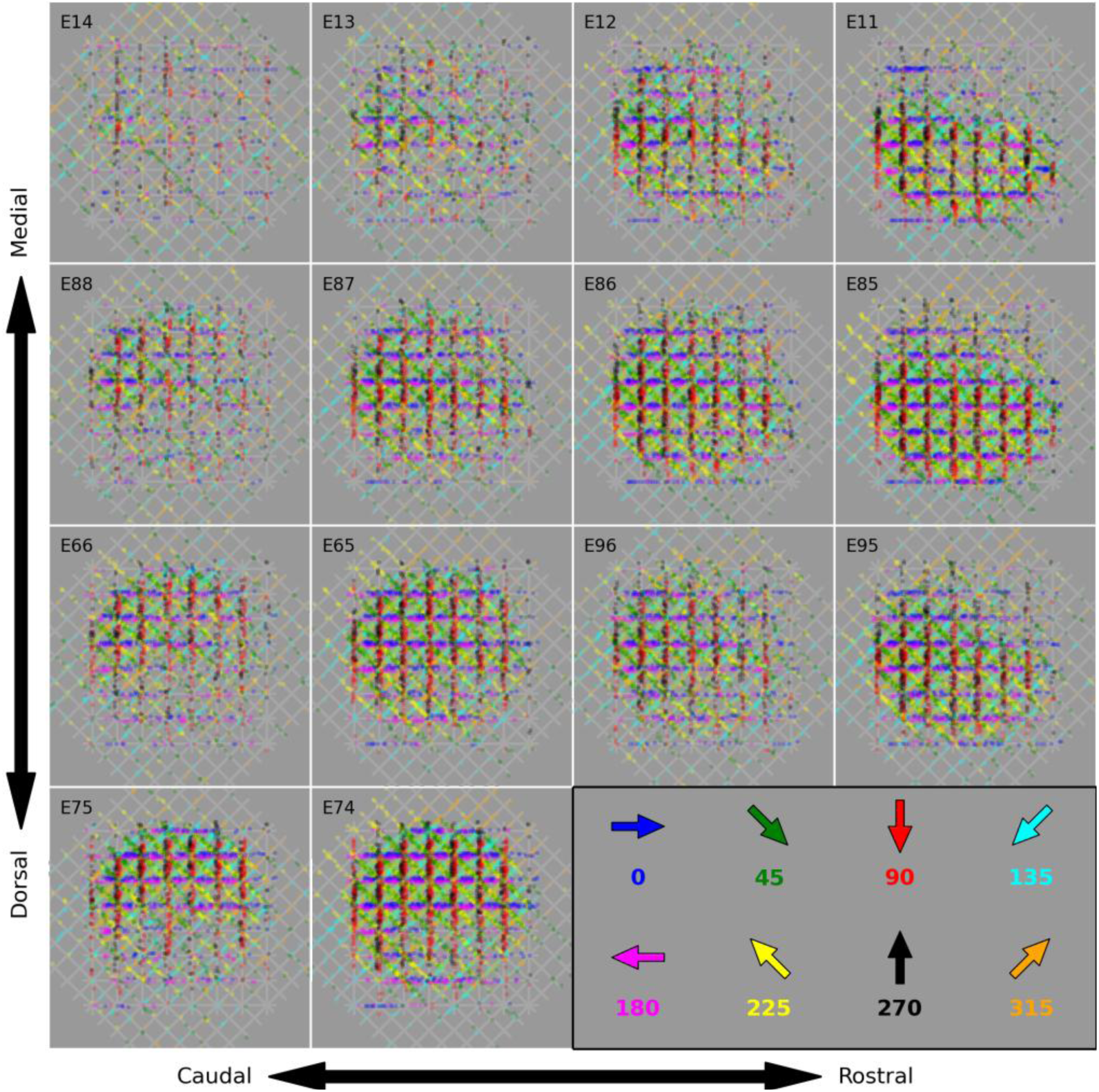
RF is large across several electrodes. RFs of 14 electrodes in a preparation are shown in response to moving dots. RFs are arranged based on the location of their electrodes in the MEA. This shows that RFs are considerably large for all the responsive electrodes. More importantly, nearby electrodes appear to have more similar RFs than distant pairs (e.g. see E74 and E75 vs. E74 and E85).

### 2. RF response similarity across the cortex

Looking at the RFs plotted across MEA, it appears that the RFs of nearby electrodes are more similar to each other than those of distant electrodes (Fig. 4 and Fig. 5a). To quantify the similarity of RFs between pairs of electrodes, we calculated the amount of overlap by binning the dots paths in the visual field to calculate the normalized average LFPs. Then we sum the smaller of the two normalized response values over all bins and divide it by the sum of the average responses over all bins (see Materials and Methods). To determine the significance of similarity between two electrodes, we recalculated the similarity between the two electrodes after shuffling the binned responses of one of the electrodes. This process was done 1,000 times. We then call the original similarity significant if it is higher than 95% of the shuffled similarities.

**Fig. 5.**
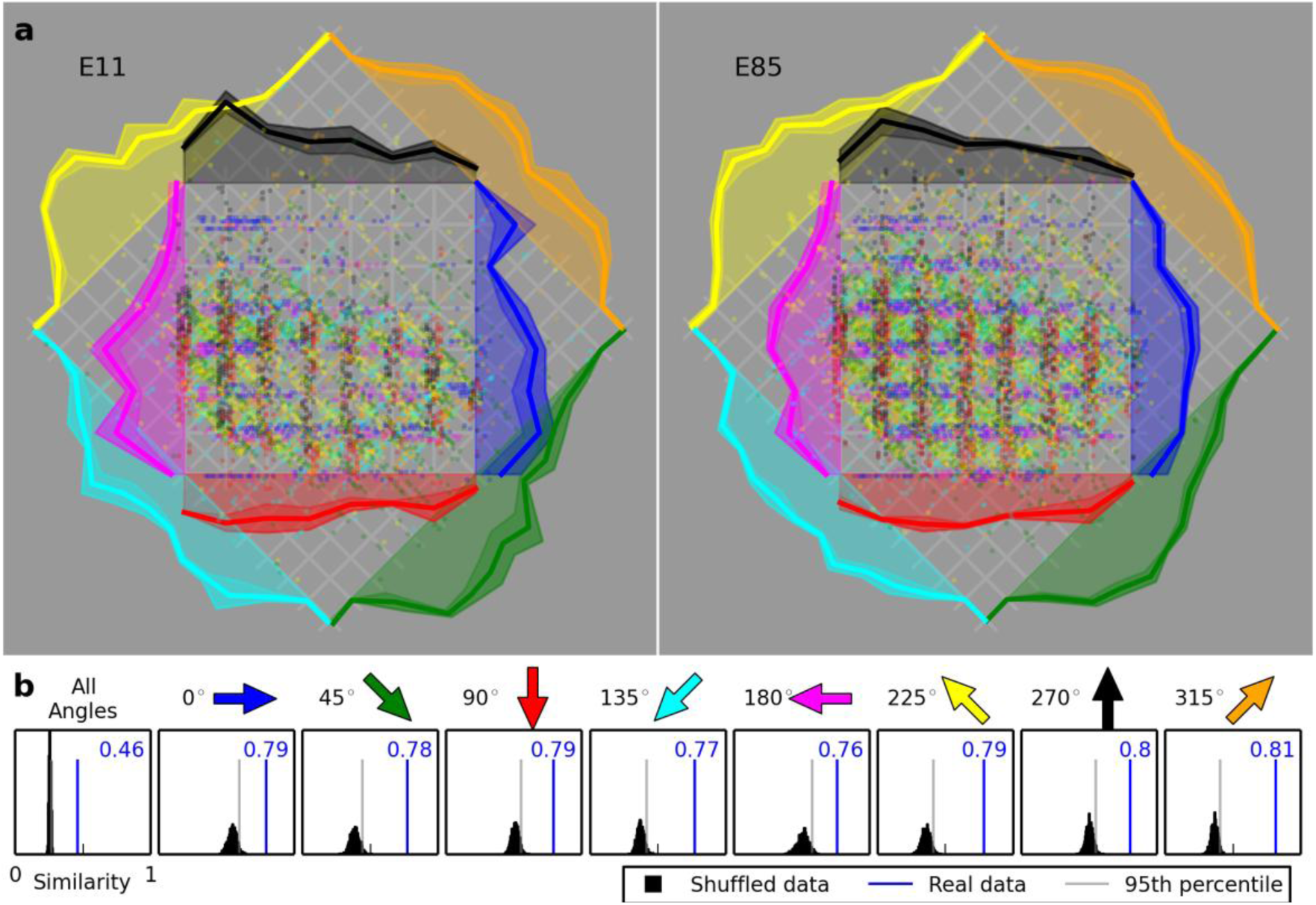
Nearby electrodes have significantly similar RFs. **a** Shown are the RFs for a nearby pair of electrodes (E85 and E11; see Fig. 4) as plotted in Fig.2. Note the similarity in the distribution of responses (solid lines) across different directions. **b** The similarity between the two RFs when the responses to dots moving at all angles are considered together (the leftmost panel), along with when we consider only the responses to dots moving at a specific angle. The black distribution shows of 1,000 calculated similarities when the data are shuffled. The light grey line shows the similarity below which 95% of the shuffled similarities lie. The blue line and number show the similarity of the real data for the two electrodes that lie well above the significant level.

Our analysis demonstrates that neighboring electrodes have significantly similar RFs, consistent with our qualitative conclusion (Fig. 5b). Regardless of the direction of moving dots, RFs of the given pair exhibit a significant similarity of 0.46. Now the question remains to be answered is that does RF similarity depend on the spatial separation between electrodes? For the four turtles that had several visually responsive electrodes, we plotted the similarity versus distance for a single electrode paired with all other electrodes. Then we made this plot for all visually responsive electrodes, and finally, we arranged these plots in the same way the corresponding electrodes are arranged on the MEA (Fig. 6a). Mostly negative slopes indicate that RFs of nearby electrode pairs tend to be more similar to each other than the RFs of distant electrode pairs. The average RF similarity at each electrode pair distance for all visually responsive electrode pairs for four turtles indicates a consistent decay (Fig. 6b).

**Fig. 6.**
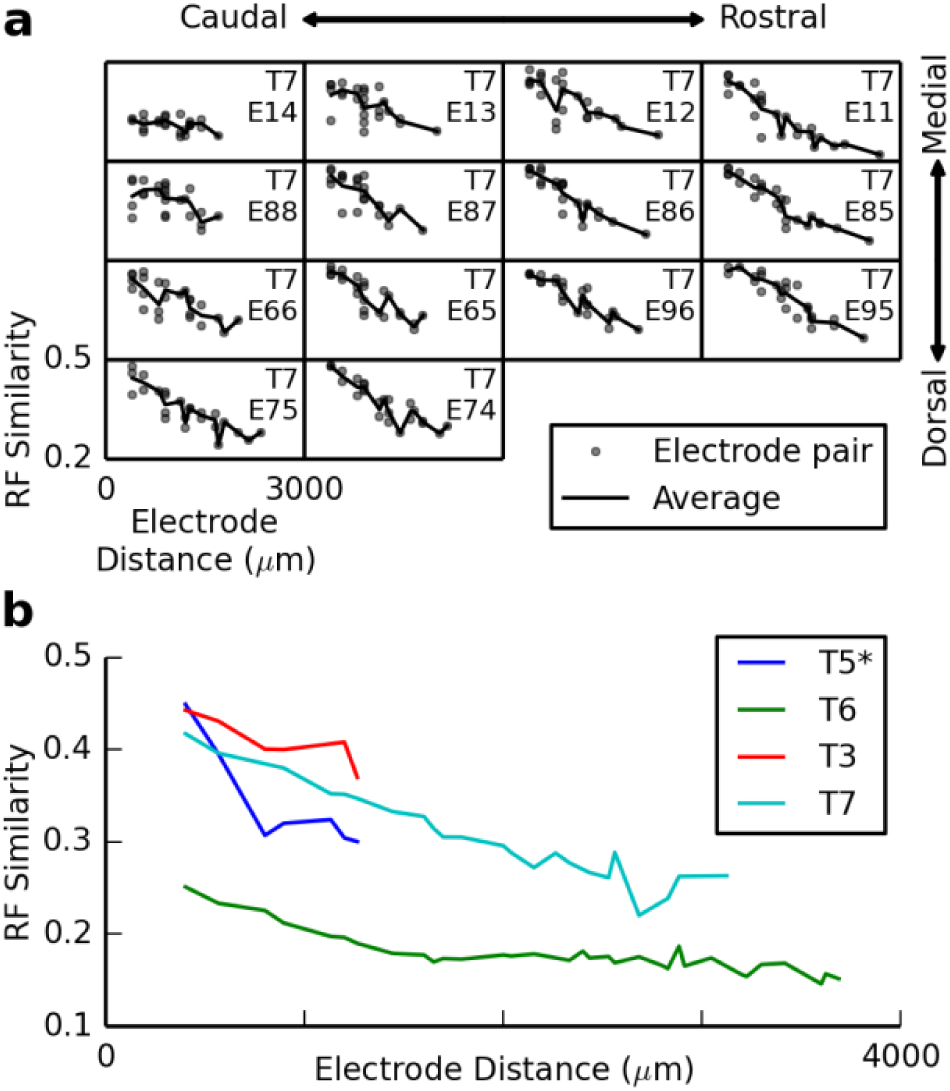
RF similarity gently decays with spatial separation between the electrode pairs. **a** Plots of RF similarity versus electrode distance. Each plot represents RF similarity of a reference electrode (show in the plot) with all other 13 electrodes (RFs are shown in Fig. 4). Plots are arranged based on the location of their reference electrode in the MEA. In each plot, at a given distance, each point is the similarity of the RF of the reference electrode with another visually responsive electrode separated with that distance. Solid line shows the average RF similarity. **b** The average RF similarity at each electrode pair distance for all visually responsive electrode pairs for four turtles. *Turtle 5 was included here to show that its trend is consistent with the others, but the visual responses for turtle 5 were relatively weak. Therefore, in order to have enough visually responsive electrode pairs for turtle 5 we used a lower threshold (0.5) for visual responsiveness.

Additionally, it appears that the negative slope can be found more consistently for the rostral electrodes than for the caudal electrodes (Fig. 6a). The caudal electrodes tend to have slopes closer to zero. This means that the RFs at caudal electrode sites are no more (or only slightly more) similar to their neighbors than they are to distant electrodes. This is seen more clearly when looking at a larger section of the array and is a consistent result across turtles (data not shown). It is worth noting that in a majority of cases; even the lower levels of similarity are still significantly more similar than shuffled data.

### 3. Direction Sensitivity

Moving dots in different directions provide us with the opportunity to look for the preference of one direction over the opposite direction (opposite directions that cover the same spatial region in the visual field). By looking at the average LFP event counts along the same path for opposite directions, we can get a sense of whether a recording site shows sensitivity to one direction compared to the opposite direction. We found no examples of an LFP being visually responsive to dots moving in one direction but not the opposite direction (Figs. 2, 3, and 5). Averages of LFP count along the paths ((solid line distributions in Figs. 2, 3, and 5) further support this hypothesis. Overwhelmingly, we discovered that the average response curves to opposite directions are nearly mirror images of each other (see Fig. 2 for example). This indicates that there is no opposite angle direction sensitivity in the LFP response (when quantifying the LFP response as the number of threshold crossing of the LFP).

### Temporal Properties

So far we have looked at the spatial structure of the responses. One equally important topic to explore is the time course of the responses. How long do responses to brief and extended stimuli last? How long is the response latency in the visual cortex? How does the strength of responses adapt? Does adaptation depend on stimulus characteristics?

LFP signals in response to brief LED flashes (50 *ms*) often contain oscillations that last for several hundreds of millisecond regardless of the flash amplitude (Fig. 7a). While at earlier times after flash onset these oscillations are clearly dominated by one or two frequencies, at later times, they are made up of fluctuations covering a broad range of frequencies (Fig. 7a). Rather than characterizing power spectrum of the responses, we quantify the temporal properties of LFP events after flash onset in terms of response duration, latency, and adaptation.

**Fig. 7.**
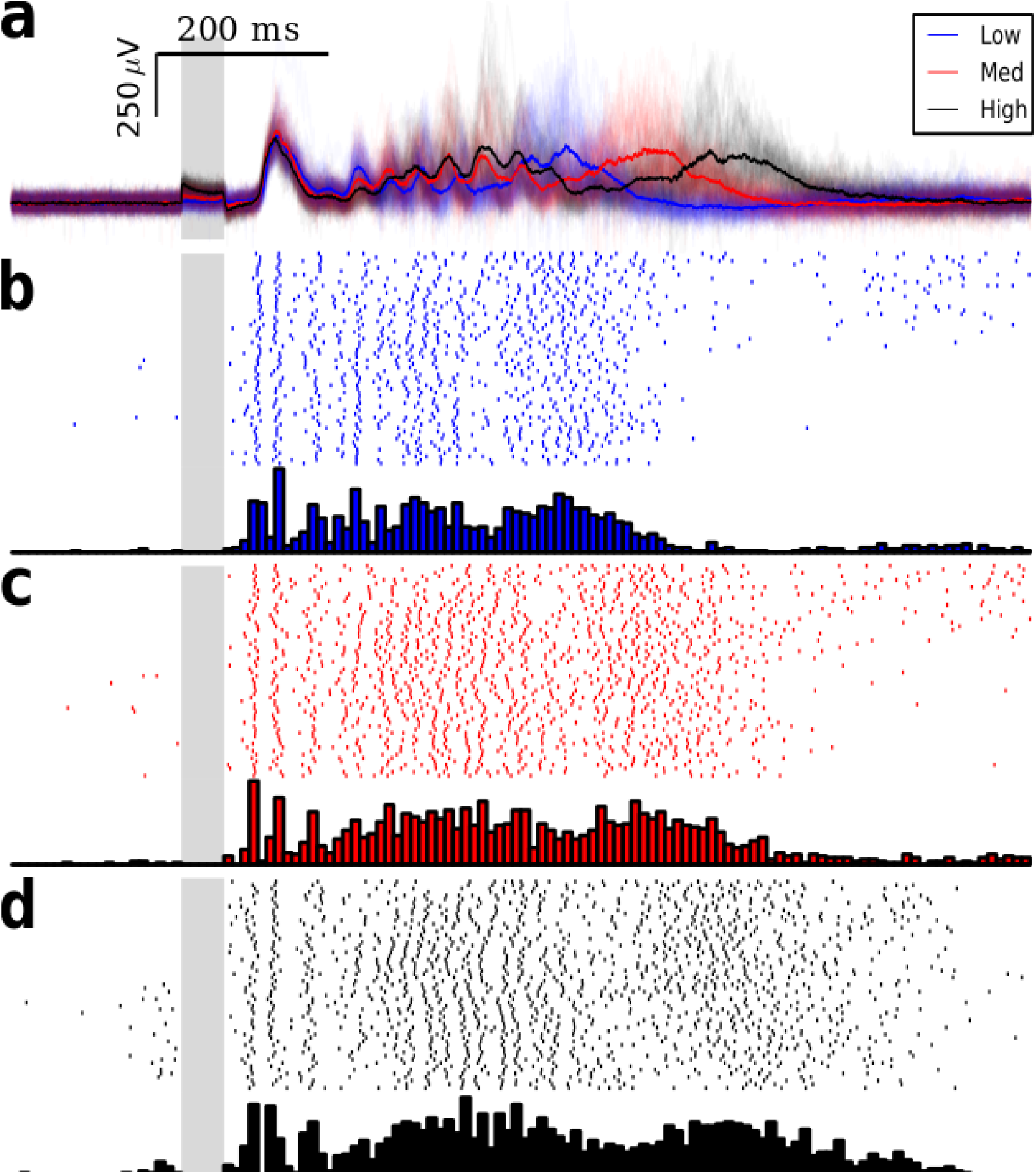
Temporal structure of the responses evoked by brief LED flashes. **a** Individual LFP responses in low opacity and average responses in bold for three amplitudes (blue, red, and black for low, medium, and high amplitudes respectively) of 50 *ms* LED flashes with 30 *s* inter-trial intervals. Early LFP responses consist of several hundred millisecond oscillations dominated by one or two frequencies while later responses cover a broad range of frequency spectrum. **b** Rastergram of LFP events (top) with prestimulus time histogram (PSTH; bottom) in response to low-amplitude flashes. Rastergram reveals the presence of oscillations in early responses. **C** and **d** Same as in **b** for medium and high flash amplitudes.

### 1. Response duration

For each trial and flash amplitude, we determine threshold crossings after stimulus onset, and then sum over trials to obtain prestimulus time histogram (PSTH; Fig. 7b, c, and d). We typically see responses that last 500 ms to 1 *s* depending on the flash amplitude. In addition, when looking at visually evoked LFP activity over an extended period of time, in many instances, we also see a second (or even third) period of activity after periods of relative inactivity even several seconds after flashes (Fig. 8). This persistent activity occurs on different electrodes of all preparations (Fig. 8) even in response to very brief flashes (10 *ms*). If we compute PSTHs for several electrodes and arrange them based on their location in MEA, we see a reliably reproducible wave of activity up to 10 s after the presentation of a flash (Fig.8).

**Fig. 8.**
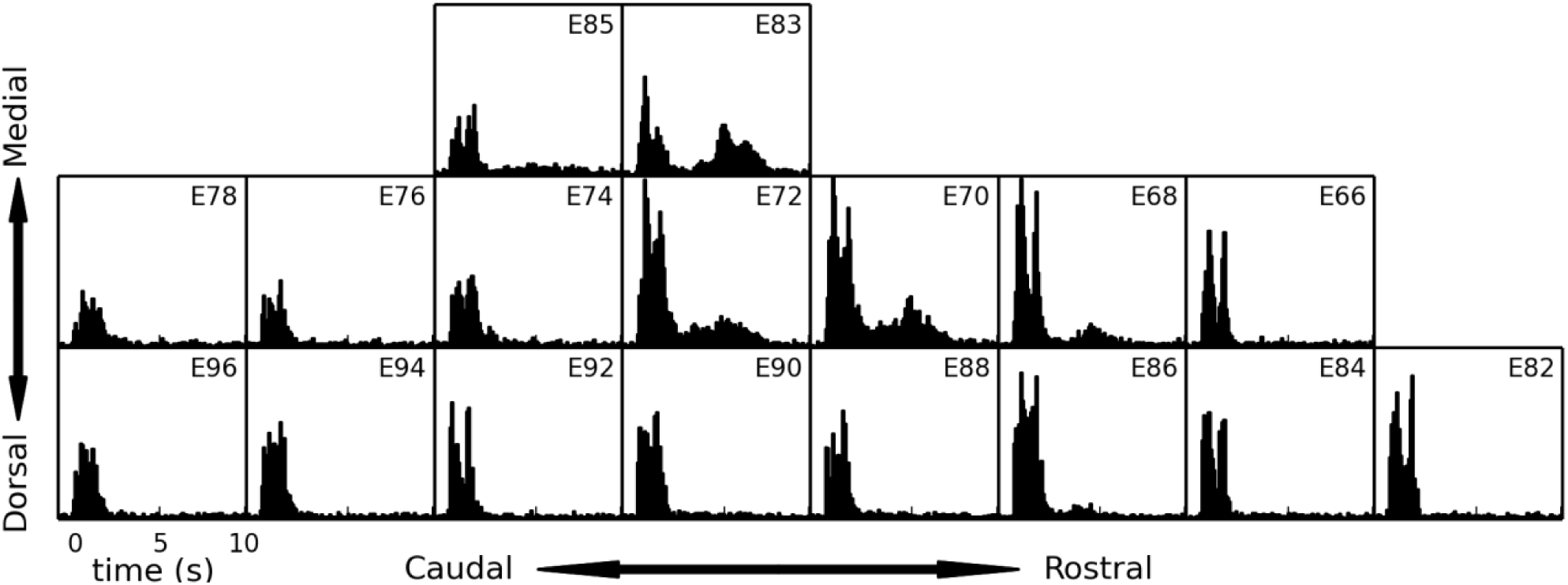
Persistent response happens on several electrodes. PSTHs are calculated over an extended period of time (10 s window) for several electrodes in response to a brief 50 *ms* LED flash. Plots are arranged based on the location of their corresponding electrodes in MEA. A second wave of activity occurs several seconds after stimulus onset and lasts for several seconds.

### 2. Response latency

Our results involve relating the activity recorded in the cortex with the stimulus presented to the retina. One important aspect of this relationship is how much delay there is between the presentation of the stimulus and the response caused by signal propagation time in the pathway leading to the cortical response. This is an interesting question per se, but also an essential piece of information when it comes to interpreting the responses to stimuli that cannot be characterized as occurring at only one instance in time.

We investigated the latency between stimulus presentation and the first evoked spike recorded on extracellular electrodes. To this end, we looked at the latency to the first spike in response to stimuli with a precise ON time covering the entire visual field. When looking at responses to a full-screen flash or to the change from a blank screen to the start of a complex movie, we found an extensively wide range of latencies (Fig. 9). Our results indicate that a typical latency to first evoked spike is around 200 - 500 *ms* (Fig. 9). A further support for this delay appears through RF analysis. For each LFP event detected, if we mark the region of the visual field in which the dot was at earlier times (0, 150, 300 *ms*), we see the most overlap of the contributions to the visual field from the dots moving at different angles (Fig. 10). As such, for all figures showing the RF as probed by moving dots, a 300 *ms* delay has been applied.

**Fig. 9.**
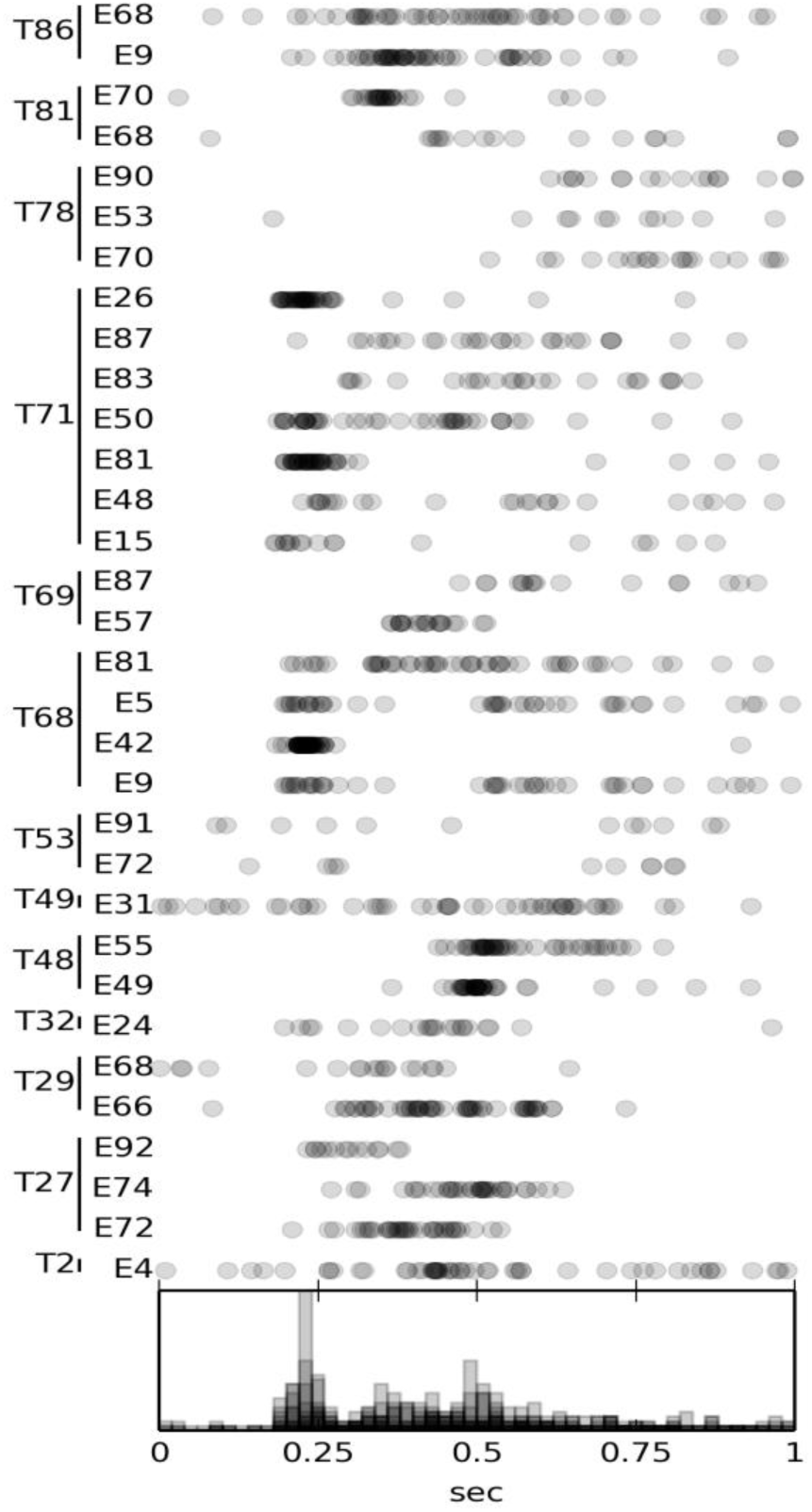
A typical latency to first evoked spike is around 200 – 500 *ms*. The time of the first detected spikes following either the onset of a red LED flash or the transition from a blank screen to the beginning of a complex movie are marked by grey circles. Latencies are shown for several trials of 32 channels across 13 different turtles. Data exhibits a significantly wide range for latency among channels, trials, and turtles. Summary histogram is shown at the bottom which shows a typical 200-500 *ms* delay. Same wavelet filtering technique is used to detect spikes, but with different parameters (Daubechies wavelets with minimum level of 3, maximum level of 7, minimum frequency of 117 *Hz*, and maximum frequency of 3750 *Hz* with a 10 times standard deviation as threshold).

**Fig. 10.**
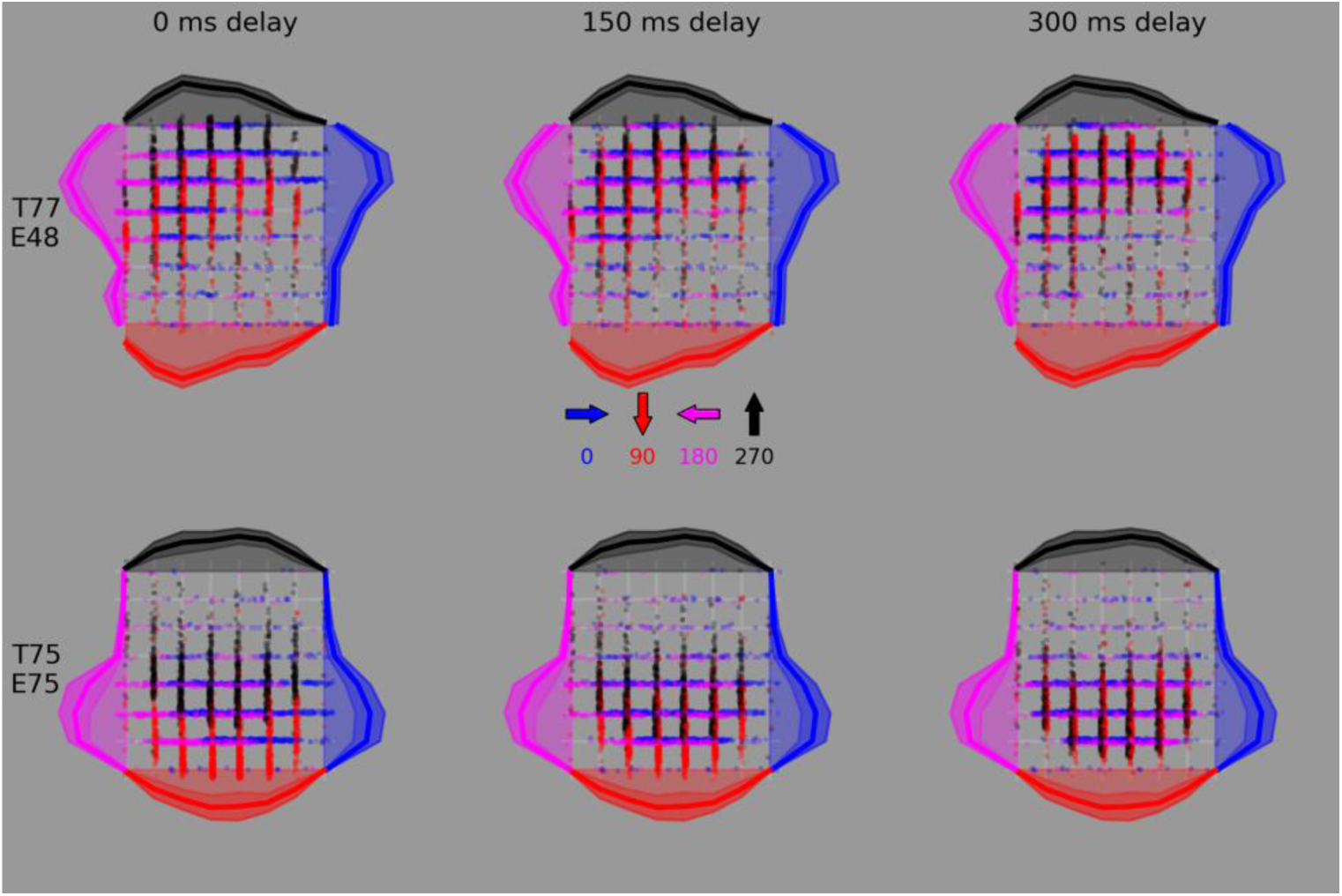
RFs converge most with delays of ∼300 *ms*. Applying a range of delays (0, 150, 300 *ms*) for two electrodes of two different turtles across several moving dot directions. Detected LFP events are attributed to the region of visual field in which the dot was at prior time. We see the most overlap of the contributions to the visual field from dots moving at different angles for a 300 *ms* time delay.

### 3. Adaptation

The effects of adaptation in turtle visual cortex are clear, long lasting, and ubiquitous. An adapted response in the visual cortex can be the result of two different sources: The cortex may be adapting in such a way that it has a diminished response (relative to the unadapted state) to the same cortical input, or adaptation has taken place at an earlier stage in the visual pathway, and the cortex is still responding in a consistent way as before adaptation but to a diminished cortical input. To determine whether cortex has an adaptation mechanism, we looked at responses in the presence or absence of prior stimulus.

#### Visual-Visual Adaptation

To test the effect of a stimulus on a subsequently presented stimulus, we used moving dots, radially moving bars, and full field flashes. In the clearest demonstration of adaptation, when we presented a series of 100 *ms* duration LED flashes to the retina, we reliably recorded a strong LFP response to the first flash, and either no response or a greatly diminished response to the subsequent flashes presented 2 *s* afterward (Fig. 11a). The extent to which the subsequent responses were diminished depended on the time separation of the flashes. Consistent with an earlier study (Luo et al., 2010), this dependence was not all-or-none; in between the short inter-flash-intervals that completely abolished subsequent responses and the long inter-flash-intervals that seemed not to affect subsequent responses, there were intermediate inter-flash-intervals that resulted in somewhat diminished subsequent responses.

**Fig. 11.**
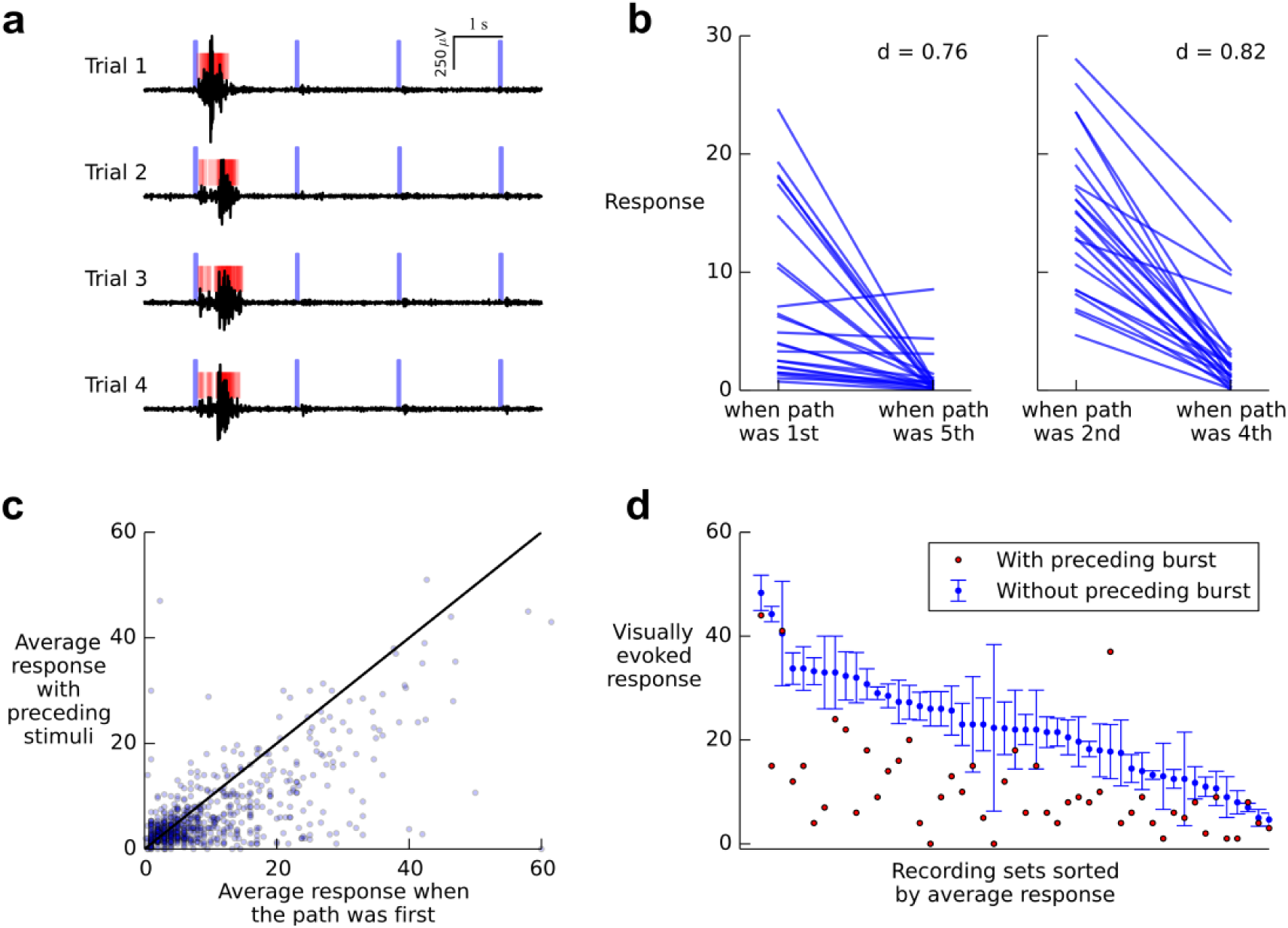
Cortical adaptation to evoked and ongoing activity. **a** LFP responses to four repeated presentations of a series of four brief full field flashes (flash timing indicated by blue vertical bars; 100 *ms* flashes with 2 *s* inter-flash-interval). LFP threshold crossings are indicated by red rasters. Adaptation abolishes responses to subsequent flashes. **b** The responses to dots moving along 5 paths across visual field greatly adapt. For each path the response strengths when the path was an early stimulus in the series of stimuli (i.e., either the first or second path to be presented) is compared against the response strengths when the same path was presented later in the series of stimuli (i.e., either the fourth or fifth path). Each point plotted is the average of 7-30 trials. The average decrease in response strength, *d*, is 0.76 (left) and 0.82 (right). **c** Average response with preceding stimuli is significantly smaller that first path average response (*p* << 0.001). **d** Response strengths with and without preceding spontaneous activity. We show visual responses from several different stimuli. Red dots indicate individual responses to visual stimuli that were preceded by a strong burst of spontaneous activity within a 5 *s* window before the stimulus. For each of those recordings, the average response of the 2-4 trials of the same stimulus nearest in time to the recording that was preceded by a burst is shown in blue with error bars showing the standard deviation. The 2-4 reference recordings were selected from recordings that were not preceded by a spontaneous burst of activity. Each recording preceded by a spontaneous burst together with the 2-4 reference recordings are collectively referred to as a recording set. Clearly spontaneous activity leads to a significant and reliable adaptation of subsequent visual responses.

We further demonstrated the effects of adaptation with more complex stimuli containing spatial and temporal structure. To quantify response adaptation to subsequent stimuli, we used dots moving along different paths through the visual field. For 8 different angles, we moved dots across 5 paths in the visual field in an ordered sequence. For opposite angles, the paths overlapped but were in reversed order, e.g. the path that was presented first at an angle of 0 degrees was presented last at 180 degrees. This allowed us to compare the response to presentation order while controlling for the area of the visual field being stimulated.

To quantify the effects of adaptation due to stimulus presentation order, we defined the decrease in response, *d*, as simply the average of the decreases for individual paths according to 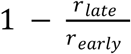, where *r*_*late*_ is the strength of the response when the path was presented late in the series (either the fifth or fourth path to be presented), and *r*_*early*_ is the strength of the response when the path was presented early in the series (either first or second).

When we look at the response amplitude when a path was presented first compared to the same path being presented fifth (or second compared to fourth), we clearly see adaptation of responses to stimulation of one area of the visual field caused by previous stimulation of other areas of the visual field (Fig. 11b).

On average, the evoked response to the fifth path is 76% smaller than when it is the first presented path. Similarly, fourth path response is 82% smaller than the second path response. The fact that the decrease in response strength is larger and more reliable for the second-fourth pairs than it is for the first-fifth pairs is most likely due to complications near the edge of the visual field. The first/fifth paths are always on the very edge of the visual field. Consequently, the dots moving along those paths were not projected to the retina nearly so long as dots moving across paths crossing a larger portion of the retina. In general, these outer paths evoked smaller responses than more interior paths. As such, these weaker responses may be more confounded by noise.

In a different set of experiments, adaptation to visual stimuli was studied while controlling for not only same path in the visual field, but also the direction of motion along that path. In contrast to the previous data set, in this data set the order in which the paths were traversed was randomized for each trial. Thus, a given path may have been the first path presented during one trial, but the fourth path presented during the next. This allowed us to separate the responses to a dot moving along any given path into trials for which the path was the first path to be presented and trials for which the path was not the first path presented. Using the same LFP threshold crossing described earlier, for each path, we calculated two average responses: the average first-presented response and the average non first-presented response. For this data set, time interval between dots moving in a given direction is 10 *s*. After presenting all 5 paths, there is either 118 *s* or 214 *s* waiting time before starting a new set of 5 dots in a different direction. In Fig. 11c the average response when a path was first is plotted against the average response to that same path when the path was not first. Because there were very few trials for any given angle-path combination, the results are somewhat scattered. But, when taken as a whole, for the 575 responsive points shown, average non first-path responses are significantly weaker than the average first-path responses (*p* = 1.7*E* - 20).

#### Ongoing-Visual Adaptation

While the results so far clearly demonstrate an adapted response to visual stimuli, they do not shed any light on the sources of adaptation. It is unclear if the adaptation is taking place in the cortex, at an earlier stage in the visual pathway, or (most likely) some combination of both effects. To better understand adaptation happening within the cortex, we looked at how visual responses adapted to spontaneous activity within the cortex (Fig. 11d). Here we looked at repeated trials of a given stimulus and picked out the trials that had a large burst of LFP events within the 5 *s* leading up to the stimulus presentation. We then calculated the average response to 4 trials of the same stimuli that did not have a large spontaneous burst preceding them. To avoid having our results confounded by experimental rundown, we selected the 4 trials that occurred most closely in time to the trial that was preceded by spontaneous activity. It is clear that spontaneous activity in the cortex can lead to a significant and reliable adaptation of subsequent visual responses (Fig. 11d). Although our results do not disentangle adaptation in the cortex from adaptation in pathways leading to the cortex, it indicates that the cortex has an adaptation mechanism by itself such that ongoing burst of activity diminishes subsequent response strength.

### Response Variability

Despite response characterization so far, we see large a variability in the responses to repeated presentations of the same stimulus. This variability manifests itself in different ways from variable strength to variable timing or even variable temporal and spectral properties (OUR PAPER).

Looking at the responses to moving dots, there seem to be two different visual responses to dots moving along the 3^rd^ and 4^th^ paths (Fig. 12). If we focus on the responses to the 3^rd^ path, we see that of the 16 trials, there are only responses in 5 or 6 of them (first of those “responses” is likely a spontaneous activity, since it starts slightly before the stimulus). On the other trials, there are no visible LFP oscillations. Two of the non-responding trials might have been affected by adaptation from the bursts of activity preceding the stimulus but that still leaves 8 non-responsive trials. Similarly, the non-responsive trials for the 4^th^ path are likely due to the responses to the 3^rd^ path that occurred just before the 4^th^path. Interestingly, this all-or-none response variability was not seen for dots moving in the opposite direction along the same paths in a set of recordings taken over the same period of time (data not shown).

**Fig. 12.**
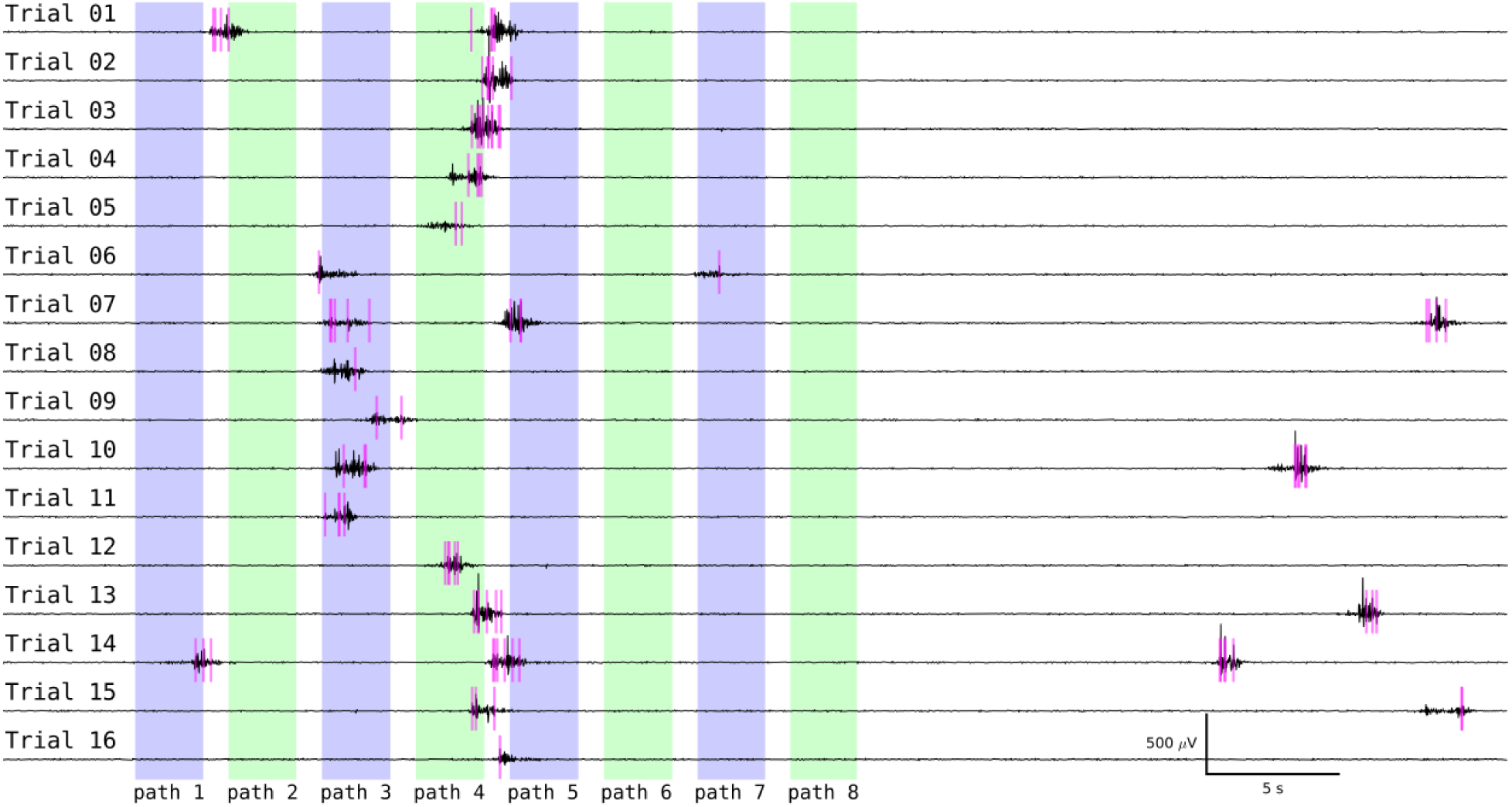
Responses exhibit variable strength, onset timing, or temporal properties. LFP signal (black) with action potentials (magenta rasters) during 16 trials of 8 dots moving along 8 different paths in one direction. The 8 colored columns indicate the timing of the 8 dots moving across the visual field. If we compare the responses to the 3^rd^ path in the 7^th^ and 8^th^ trial, we find markedly different response amplitudes. Response onset time to the 4^th^ path shows variability of order of several seconds as well as in how that response plays out. For some trials (e.g., trials 1-3) we see roughly one large burst, and for others (e.g., trial 4) it looks more like a series of two smaller bursts.

We also see variability in the temporal structure of the response (Fig. 12). Not only do responses to both the 3^rd^ and the 4^th^ path have substantial differences in the time to response onset (sometimes varying by as much as a second), but they also vary in how that response plays out. For some trials (e.g., trials 1-3) we see roughly one large oscillation, and for others (e.g., trial 4) it looks more like a series of two smaller bursts.

Finally, Fig.12 also contains examples of response strength variability. If we compare the responses to the 3^rd^ path in the 7^th^ and 8^th^ trial, we find markedly different amplitudes of response. These interesting observations worth exploring further in future studies.

### Spike-LFP Correlation

In general, spikes are much less common in the absence of LFP activity than they are during a burst of LFP activity and there is a clear positive correlation between the number of action potential and the number of LFP peaks (Shew et al., 2015). Here we look at the RF size using detected spikes and then compute its similarity to the RF obtained using LFP events. Again, spike RFs cover a large region of the visual field and, in other words, units fire action potentials in response to dots presented at different locations of the visual field (Fig. 13a). However, spikes have a less reliable trial-to-trial variability since they are less common than LFP events. More importantly, spike RF is exceptionally (∼80%) similar to the RF obtained using detected LFP events on the same electrode (Fig. 13b; average overall similarity 45 ± 15%, *mean* ± *std*).

**Fig. 13.**
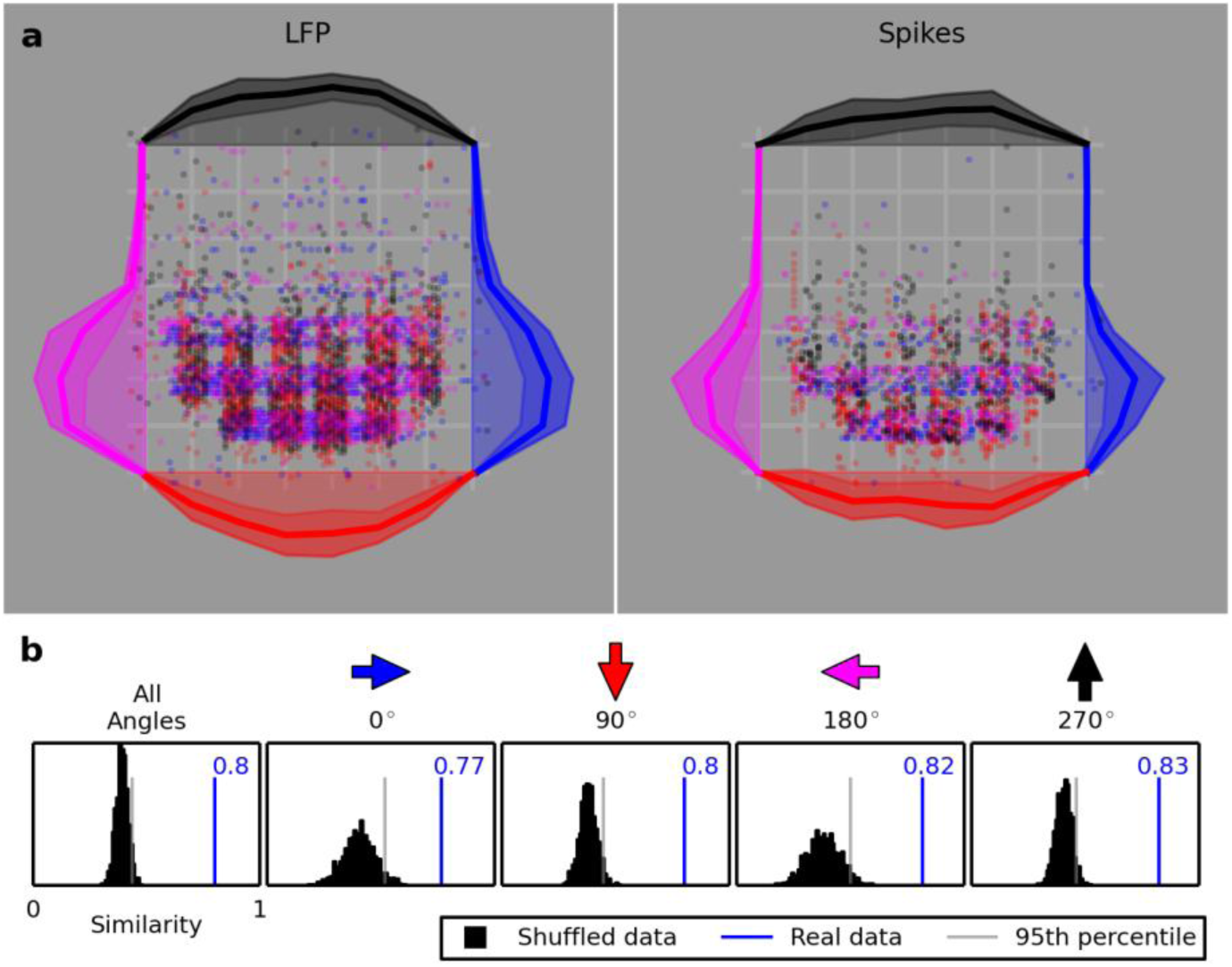
Spike-LFP RF exhibits a significant similarity. **a** The LFP (left) and spike (right) RFs for one electrode are plotted. The spike RF is obviously smaller that the LFP RF but still considerably large. **b** The similarity between the two RFs when the responses to dots moving at all angles are considered together, along with the similarity separately for different angles. The black distribution shows of 1,000 calculated similarities when the data are shuffled. The blue line and number show the similarity of the real data, and the light grey line shows the similarity below which 95% of the shuffled similarities lie.

### Possible Mapping of the Visual Field to the Visual Cortex

For decades it has been reported that pyramids in turtle visual cortex respond to small moving stimuli spanning a very large portion of the visual field. While our results show large RFs for both LFP events and spiking activity, when we looked for more detail in the spatial structure of the RFs, there is less of a consensus regarding what to expect. Though there is not a clear and well-defined retinotopic map to the cortex, there have been a few studies that report on projection at two steps leading from the retina to the cortex. It has been hypothesized that naso-temporal axis of visual space is represented along the rostro-caudal axis of the visual field (Mulligan and Ulinski, 1990). However, this prediction contradicts earlier results indicating opposite polarity in recorded evoked potentials in response to localized visual stimulation of the retina (Mazurskaya, 1973a).

To resolve this puzzle, we used moving dot stimuli and investigated whether the nasal-temporal response specificity changes as we compare data from electrodes in the rostral cortex with those from caudal cortex. By performing the same experiments with the visual stimuli rotated 90 degrees, we were able to investigate dorso-ventral response specificity. We found that neither Mazurskaya’s results nor Mulligan and Ulinski’s predictions are consistent with our findings.

#### Naso-temporal Visual Field

We tested this with moving dots that followed straight paths from the top of the visual field to the bottom (as well as dots moving in the opposite direction). Eight of these vertical paths were spread out at different naso-temporal locations spanning the visual field. Only one dot (following one path) would move at a time. After moving these dots along the different paths we could look for naso-temporal response specificity at any given recording site in the cortex to see if the nasal-temporal response specificity changes as we compare data from electrodes in the rostral cortex with those from the caudal cortex.

Our results show that, as you compare the naso-temporal response specificity of different recording sites, the strength of response specificity does change, but the pattern of specificity was approximately the same for all recording sites, and the variations from site to site did not follow any clear trend (e.g., rostral recording sites responding strongly to one area while caudal sites respond strongly to a different area; Fig. 14a).

**Fig. 14.**
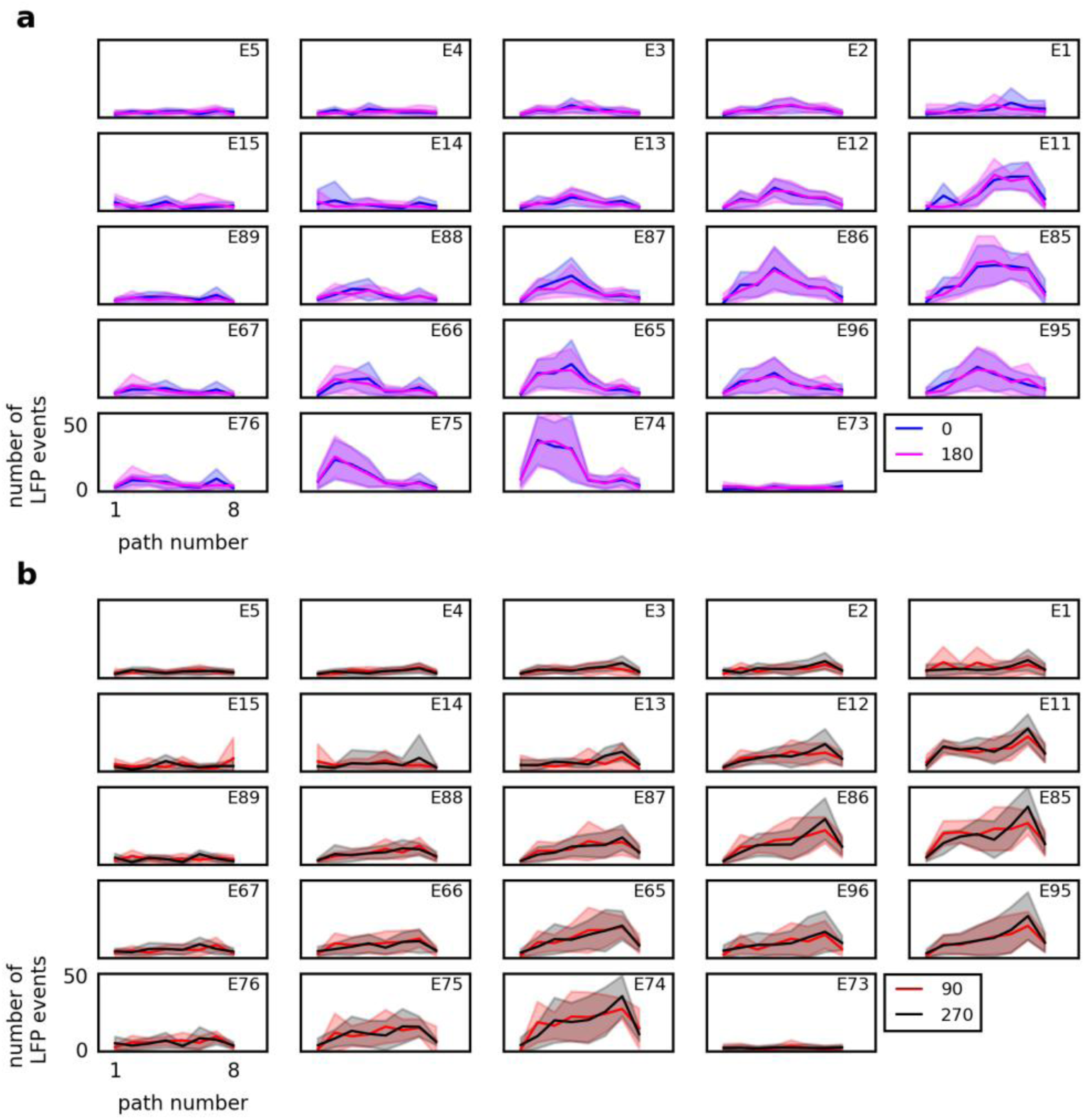
No clear mapping from the visual field to the visual cortex exist. **a** Naso-temporal response specificity has variable amplitude but similar patterns for the recording sites on MEA. For 24 electrodes we show the average responses (lines) and standard deviations (filled area around the lines), in response to dots moving along 8 different vertical paths arranged naso-temporally in the visual field. The two colors represent the two opposite angles that traverse the paths. Response strength varies but not the pattern and, moreover, the variations from site to site did not follow any clear trend. **b** LFP responses to dots moving along 8 paths arranged vertically in the visual field plotted in the same format as in **a**. The two colors represent the two opposite angles.

#### Dorso-Ventral Visual Field

By performing the same experiments with the visual stimuli rotated 90 degrees, we investigated response specificity for different elevations in the visual field. In contrast with previous predictions, we found recording sites that responded clearly to only the upper or only the lower visual field rather than stimulation at all elevations (Fig. 14b). Overall these findings strongly support no well-defined mapping from the visual field to the visual cortex.

## DISCUSSION

Here, we investigated the basic spatiotemporal properties using MEA recordings of visual responses in turtle whole-brain eye-attached *ex-vivo* preparations while stimulating retina primarily with black moving dots in different directions. We found large RFs that typically cover over half of the visual field with no indication of direction sensitivity. Our investigations revealed no clear or well-defined retinotopic map from the visual field to the cortex. Additionally, a broad range of response latencies and a strong adaptation to both ongoing and visually evoked activity have been observed.

We know that the size of RFs in lateral geniculate nucleus (LGN) cells is restricted to 30 degrees (Boiko, 1980) and that there are thalamic axons connecting to the cortex with RFs as small as 2-5 degrees (Mazurskaya, 1973a). Then how does large cortical RF settle with small RF found in the LGN? Two explanations are conceivable here. It could be that an individual cortical cell has a large RF because it receives input from many LGN cells whose RFs collectively span a large area of the visual field, but these seems unlikely given the proposed projections from LGN to cortex (Mulligan and Ulinski, 1990). Therefore, what seems more likely is that the large cortical RFs may be the result of individual cortical cells receiving input from many other cortical cells that each receives LGN input representing only a small portion of the visual field.

While it is important to understand the thalamic inputs to the cortex, we firmly believe that intracortical connections play an essential role in determining the RF size and structure. The extent to which the RF of individual cells in the visual cortex depends on the connectivity with other cortical cells has been demonstrated by comparing the normal RF of a cortical cell to its RF after applying pharmacological blockers to different areas of the cortex (Mazurskaya, 1973b). In support of our viewpoint, this study found that after applying blockers to other areas of the cortex, there would be gaps in the large RF that previously were not present. This suggests that for that cell, its responsiveness to certain regions of the visual field depended on receiving signals from the blocked region of the cortex.

How the RF similarity of nearby pairs can be interpreted? It might suggest that LFP reach is larger than the inter-electrode distance (∼400 μm) so that neighboring electrodes are essentially measuring the same signal. While there have been some recent claims that the spatial extent of the LFP can be as large as several millimeters (Kajikawa and Schroeder, 2011), it has usually been thought that the LFP represents neural activity within roughly 150 - 400 μm of the electrode (Katzner et al., 2009; Xing et al., 2009). The fact that we occasionally see bursts in narrow frequency bands on one electrode but not on the adjacent electrode (data not shown) is consistent with a smaller spatial extent for the LFP. This suggests that the similar RFs recorded at different electrodes (spaced 400 μm apart) are not merely measurements of the same signal generated by common sources, but are instead measurements of activity generated by different sources. A reasonable speculation is that they happen to produce similar signals due to immense recurrent connections among neurons.

Our findings overwhelmingly point to no direction sensitivity in moving dot experiments and this is in agreement with previous studies. It has been reported that only 9% of units in turtle thalamus show direction sensitivity, though the significance of these results is questionable (Boiko, 1980). Whether or not direction sensitivity exists in the thalamus affects how we think about its occurrence in the cortex. In either case, we have to take synaptic connection structure into account to explain direction insensitivity of cortical neurons to given thalamic inputs.

It takes time for the signal elicited by visual stimuli to travel from the retina to the thalamus and then to the cortex. Many studies have reported first spike latencies in visual cortex to be between 80 ms and 200 ms (Mazurskaya, 1973a) and even as short as 25 - 150 ms (Bass et al., 1983), and latency to LFP response onset of 86 ± 4 ms (Prechtl and Bullock, 1994). However, we found that, in responses to a full screen flash or to the change from a blank screen to the start of a complex movie, a typical latency to first spike is around 200 - 500 *ms* which is longer than previously reported latencies. In addition, we provided further support for this range of time delays by looking at the structure of RFs and making it maximally convergent.

The effects of adaptation in turtle visual cortex have long been studied. Our investigation stated that the cortex has an adaptation mechanism but did not speak about adaptation in neural pathways leading to the cortex. It is almost certainly the case that both of these effects contribute to the adaptation observed in the cortex. Adaptation in our preparations seems to persist for several seconds to a minute in line with others that showed complete recovery happens in ∼16 s (Luo et al., 2010). Though some studies described recovery times in visual cortex ranging from 0.5 min to 3 min (Gusel’nikov et al., 1972). We suspect that the apparent discrepancy is due to the different stimuli they used.

Variability we observed in turtle visual cortex has also been seen in cat visual cortex in voltage sensitive dye recordings (Holt et al., 1996; Sadagopan and Ferster, 2012). It has been shown that much of the trial-to-trial variability could actually be explained by the ongoing activity in the cortex (Arieli et al., 1996). That is to say that after subtracting the activity of the cortex immediately preceding the response, the variability of the responses was greatly reduced. More generally, it has been suggested that sensory responses should be thought of as not simply the product of a sensory input and some “default” anatomical connectivity, but instead the product of those along with learned expectations and environmental contingencies that can change continuously (Fontanini and Katz, 2008).

A complication our results point to is the presence of a well-defined retinotopic map from the visual field to the cortex. Using retinal ablation and observing orthograde degeneration, Ulinski and Nautiyal (1988) reported that the nasal retina projects to the contralateral rostral lateral geniculate nucleus (LGN). Later, Mulligan and Ulinski (1990) found that the rostral LGN projects to the caudal cortex and the caudal LGN projects to the rostral cortex. Combining these two observations they predicted that the nasal-temporal axis of visual space is represented along the rostro-caudal axis of the visual cortex. This prediction contradicted earlier results from Mazurskaya(1973), who observed the opposite polarity in recorded evoked potentials in the visual cortex while presenting local visual stimulation to the retina.

Ulinski and Nautiyal suggest that the dorso-ventral axis of the retina projects along the dorso-ventral axis of the LGN, but the data supporting this claim were much less clear than the data supporting conclusions about naso-temporal projections. Continuing along this visual pathway, Mulligan and Ulinski reported that (at least some) neurons in any given dorso-ventral transect of the LGN project along the full lateral-medial extent of the cortex. Thus, a neuron located anywhere along a lateral-medial line in the cortex can respond to stimulation at any point along a particular vertical line in the visual field. With moving dot stimuli, we found that response specificity has variable amplitude but similar patterns for different recording sites. Also, response variations from site to site did not follow any clear trend. We are convinced that experimental data supports our conclusions better than previous studies.

What do we learn by comparing visual cortex in turtles with mammals? Inferior temporal cortex (IT) is a visually responsive area in mammals that has a striking resemblance to what we found in turtle cortex. In IT responses to visual stimuli persist up to 15 *s* (Fuster and Jervey, 1981) and its cells also tend to have large RFs, respond to many stimuli, including moving stimuli, and have adaptation effects with inter-stimulus intervals less than 5 *s* (Gross et al., 1972). Unlike turtle visual cortex strong direction sensitivity has been observed in mammalian IT. Some of the direction sensitive IT cells had one clear preferred direction (termed unidirectional), but most were bidirectional sensitive (they responded preferentially to both a direction and the opposite direction, but not perpendicular motion). This was demonstrated in IT using black bars sweeping across the visual field while we have used black dots that take up only a small portion of the visual field. To scrutinize the similarity, further studies should be done with moving bars that span the entire visual field.

Mammalian hippocampus and piriform cortex are other areas that have a similar structure to turtle cortex. Like turtle visual cortex, the mammalian hippocampus has extensive feedback connections to its primary input source (entorhinal cortex; Witter, 1993). In the hippocampus, like turtle cortex, oscillations are found in the gamma band (Bragin et al., 1995) and theta band with electrodes spanning several hundred microns having similar LFP signals (Buzsáki, 2002).

The olfactory or piriform cortex is a three layer cortical structure, and has feedforward and feedback circuits that are similar to those found in turtle dorsal cortex along with numerous other similarities (Haberly, 1985). Further identifying structural and functional similarities between the turtle dorsal cortex and mammalian piriform cortex will likely help elucidate common organizational and computational principal of cortical networks (Fournier et al., 2014).

Though we have not yet seen the multitude of studies on turtles as we’ve seen with other preparations such as rat, mouse, and cat; the turtle preparation is becoming more appreciated for allowing the experimenter to study cortical processing from the subcellular level to the level of neuronal ensembles simultaneously, as well as being tolerant enough to a wide range of flexible modifications to meet the needs of a range of experiments with different technical demands.

## ACKNOWLEDGEMENTS

We thank members of the Neurophysics Laboratory for helpful discussions.

### CONFLICT OF INTEREST STATEMENT

No conflict of interest exists for any of the authors.

### ROLE OF AUTHORS

JP and RW devised the study; JP, NW, WC, WS, and RW performed the experiment; MH and JP analyzed the data; MH, JP, and RW wrote and revised the manuscript.

